# Post-hoc modification of linear models: combining machine learning with domain information to make solid inferences from noisy data

**DOI:** 10.1101/518662

**Authors:** Marijn van Vliet, Riitta Salmelin

## Abstract

Linear machine learning models “learn” a data transformation by being exposed to examples of input with the desired output, forming the basis for a variety of powerful techniques for analyzing neuroimaging data. However, their ability to learn the desired transformation is limited by the quality and size of the example dataset, which in neuroimaging studies is often notoriously noisy and small. In these cases, it is desirable to fine-tune the learned linear model using domain information beyond the example dataset. To this end, we present a framework that decomposes the weight matrix of a fitted linear model into three subcomponents: the data covariance, the identified signal of interest, and a normalizer. Inspecting these subcomponents in isolation provides an intuitive way to inspect the inner workings of the model and assess its strengths and weaknesses. Furthermore, the three subcomponents may be altered, which provides a straightforward way to inject prior information and impose additional constraints. We refer to this process as “post-hoc modification” of a model and demonstrate how it can be used to achieve precise control over which aspects of the model are fitted to the data through machine learning and which are determined through domain information. As an example use case, we decode the associative strength between words from electroencephalography (EEG) reading data. Our results show how the decoding accuracy of two example linear models (ridge regression and logistic regression) can be boosted by incorporating information about the spatio-temporal nature of the data, domain information about the N400 evoked potential and data from other participants.

**Highlights:**

- We present a framework to decompose any linear model into three subcomponents that are straightforward to interpret.
- By modifying the subcomponents before re-assembling them into a linear model, prior information and further constraints may be injected into the model.
- As an example, we boost the performance of a linear regressor and classifier by injecting knowledge about the spatio-temporal nature of the data, the N400 evoked potential and data from other participants.

## 1 Introduction

Linear models are the workhorse behind many of the multivariate analysis techniques that are used to process neuroimaging data,^1^ with applications ranging from signal decomposition^2 3 4^ to source modeling^5 6 7 8^ and signal decoding.^9 10 11^ Even though they may serve very different purposes, the data transformation performed by all linear techniques can be mathematically described by a single matrix multiplication between the input data and a “weight matrix”. From this point of view, the key difference between the various techniques is how the weight matrix is computed.

Supervised linear machine learning algorithms compute the weight matrix based on examples of the input data and the desired output.^12^ This class of algorithms have advanced the analysis of neuroimaging data on two important fronts. First, by learning what is signal and what is noise, the signal can be projected away from noise sources, which provides an alternative method to increase signal-to-noise ratio (SNR) to signal averaging. This makes it for example possible to perform single-subject and even single-trial analysis.^13 14 15^ Second, by focusing on patterns rather than individual data points, there is no longer a requirement for a one-to-one correspondence between the experimental manipulation and a change in the signal at a certain location, time, or frequency, which enables more ambitious neuroimaging studies.^16 17^

The success of machine learning algorithms to find the desired transformation is for a large part dependent on the ratio between the number of parameters that need to be estimated and the number of provided training examples. In general, the more parameters that need to be estimated, the more training data is needed to prevent overfitting of the model.^18 19^ Unfortunately, it is common in neuroimaging studies for the data dimensionality to exceed the number of trials in a recording, in which case restrictions need to be placed on the model in order to force a unique solution. Especially in these cases, it is desirable to inspect the data transformation that was “learned” by the algorithm to understand what aspects of the data contribute to the output of the model, identify possible problems, and possibly impose further restrictions on the model if the transformation was unsatisfactory.

In linear models, there are some effective general purpose approaches to place restrictions on the learned data transformation, notably *ℓ*_1_ regularization,^20^ which enforces sparsity of the weight matrix, and *ℓ*_2_ regularization,^21^ which enforces a small magnitude of the individual weights. Moving beyond these approaches, imposing further restrictions that are motivated by domain information may lead to even better performance of the model. However, it is in practice very difficult to express domain information in terms of the weight matrix,^22^ since interpreting this matrix is not straightforward when there are co-linearities in the data, which is almost always the case in neuroimaging.

To facilitate the interpretation of linear models, Haufe et al. (2014) introduced a way to transform the weight matrix into a pattern matrix, which is easier to interpret (see section 2.2). While Haufe et al. (2014) focused on the computation, visualization and interpretation of the pattern matrix, they suggest that their work may have applications stretching beyond model interpretability and form the basis for a method for incorporating domain information into linear models. In the current paper, we continue this line of thought, leading to what we call the “post-hoc modification” framework.

It is often more straightforward to formulate domain information in terms of the pattern matrix than the model weights. This has been long known in the domain of electrophysiological source estimation of electroencephalography (EEG) and magnetoencephalography (MEG) data, where the pattern matrix corresponds to the leadfield (i.e., forward solution) and the weight matrix to the inverse solution. Methods for estimating EEG/MEG source activity often formulate their domain information driven priors on the leadfield.^23 24 25 26^ The modified leadfield is afterwards combined with a sensor-to-sensor covariance matrix and inverted to yield an inverse model that incorporates the domain information. In this paper, we combine the insight of Haufe et al. (2014) that a pattern matrix can be computed for any linear model, with the insight from source estimation methods that priors that are formulated on the pattern matrix can be translated into priors on the weight matrix.

In our framework, we decompose the weight matrix of a linear model into three subcomponents, and hence divide the problem of estimating the weight matrix into three subproblems (see section 2.2):

1. the pattern matrix of Haufe et al. (2014), associated with the subproblem of determining signal components that carry information about the desired output
2. the data covariance, associated with the subproblem of estimating the relationships between model inputs
3. the normalizer, associated with the subproblem of fine-tuning the mapping between the model output and the desired output

Inspecting these subcomponents in isolation offers an intuitive way to gain insights into the functioning of the model and possible problem points. We then proceed by modifying each component to impose new constraints and incorporate domain information, before recomposing the subcomponents back into a weight matrix. Since the decomposition-modification-recomposition cycle of the weight matrix takes place after the initial model has been constructed through a conventional machine learning algorithm, we refer to this process as “post-hoc modification”.

While the framework is agnostic to the methods by which the initial linear model was constructed, and is hence applicable to a wide variety of data analysis techniques, we will use linear regression as an example throughout this paper to provide context to our procedures and equations. To provide practical examples, we demonstrate several ways in which the framework may be used to combine machine learning with domain information to decode the associative strength between words from an EEG recording, following a semantic priming protocol.^27 28^ We explore a regression scenario with a ridge regressor as a base model, and also a classification scenario with a logistic regressor. Using the post-hoc modification framework, these two general purpose models were modified to incorporate 1) the dependency between EEG sensors and time samples, 2) data recorded from the other participants, and 3) the timing of the N400 component of the event-related potential (ERP), which occurs around 400 ms after the onset of the second word stimulus.^29 30^

## 2 Methods

### 2.1 Linear models

The post-hoc modification framework can operate on any type of linear model, regardless of function and type of data, so there are many application areas. Since our examples are in the domain of machine learning, we have chosen to adopt the general purpose terminology used in that literature^31^ See Table 1 for a summary of the mathematical symbols used in this paper.

**Table 1:** A summary of the mathematical symbols used in this paper.

|  |  |
| --- | --- |
| $k$ | Number of targets |
| $m$ | Number of features describing an observation |
| $n$ | Number of observations in a dataset |
| $\mathbf{x}$ | A row vector of length $m$ that describes a single observation |
| $\mathbf{X}$ | A dataset consisting of $n$ observations |
| $\hat{\mathbf{X}}$ | An approximation of $\mathbf{X}$ |
| $\Sigma_{\mathbf{X}}$ | The empirical covariance matrix of $\mathbf{X}$ |
| $\tilde{\Sigma}_{\mathbf{X}}$ | A modified version of the empirical covariance matrix of $\mathbf{X}$ |
| $\mathbf{y}$ | A row vector of length $k$ that describes the desired output for a single example observation |
| $\mathbf{Y}$ | The desired output of a model for $n$ example observations |
| $\hat{\mathbf{Y}}$ | The actual output of a model, here an approximation of $\mathbf{Y}$ |
| $\Sigma_{\hat{\mathbf{Y}}}$ | The empirical covariance matrix of $\hat{\mathbf{Y}}$, also referred to as the normalizer |
| $\tilde{\Sigma}_{\hat{\mathbf{Y}}}$ | A modified normalizer |
| $\mathbf{W}$ | The weight matrix describing a linear transformation from $\mathbf{X}$ to $\hat{\mathbf{Y}}$ |
| $\tilde{\mathbf{W}}$ | The updated weight matrix obtained by combining $\tilde{\Sigma}_{\mathbf{X}}$, $\tilde{\mathbf{P}}$ and $\tilde{\Sigma}_{\hat{\mathbf{Y}}}$ |
| $\mathbf{P}$ | The pattern matrix describing a linear transformation from $\hat{\mathbf{Y}}$ to $\hat{\mathbf{X}}$ |
| $\tilde{\mathbf{P}}$ | A modified pattern matrix |
| $\mathbf{I}$ | An identity matrix of appropriate size |
| $\lambda$ | Controls the amount of $\ell_2$ regularization of the covariance matrix |
| $\alpha$ | Controls the shrinkage of the spatial component of the covariance matrix |
| $\beta$ | Controls the shrinkage of the temporal component of the covariance matrix |
| $\mu$ | Controls the center of the Gaussian kernel used as a windowing function for the pattern matrix |
| $\sigma$ | Controls the width of the Gaussian kernel used as a windowing function for the pattern matrix |
| $\rho$ | Controls the weighting between the pattern matrix for the current recording and the grand-average pattern matrix across all other recordings |

We will refer to a data instance, for example a single epoch of EEG data or a single functional magnetic resonance imaging (fMRI) image, as an “observation”. An observation consists of *m* “features”, for example the voltage at each sensor and each each time point of an epoch, or the beta weight for each voxel in an fMRI image. In this manner, a single observation is described by row vector **x** ∈ ℝ^1×*m*^ and an entire data set, consisting of *n* observations, by matrix **X** ∈ ℝ^*n*×*m*^.

A linear model transforms the input data by making a linear combination of the *m* features to produce output data with *k* dimensions, referred to as “targets”. In machine learning, the desired transformation is deduced by exposing the algorithm to an example input data set **X** along with the desired output **Y** ∈ ℝ^*n*×*k*^. This process is referred to as “training” the model.

To simplify the equations, it is assumed, without loss of generalization, that the columns of both **X** and **Y** have zero mean. In practice, this can be achieved by removing the column-wise mean from **X** and **Y** before entering them into the model and adding the removed offsets back to the output. Under the zero-mean assumption, the data transformation that is performed by a linear model can be represented by a multiplication between **X** and a weight matrix **W** ∈ ℝ^*m*×*k*^:

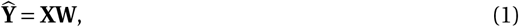

where 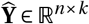 denotes the output of the model. In the case of machine learning, **W** is chosen such that 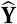 approximates **Y**, given a certain data-fit cost function (also known as a loss function). Example cost functions are the sum of squared errors, often used in linear regression, and the logistic loss function in the case of logistic regression.

### 2.2 Post-hoc modification

Haufe et al. (2014) showed the relationship between a linear decoding model **W** that approximates **Y** given **X** and the corresponding encoding model **P** ∈ ℝ^*m*×*k*^ that does the opposite and approximates **X** given 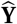:

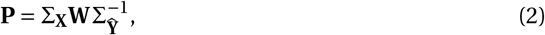

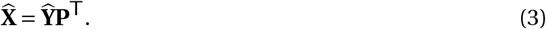

In the above equations, 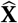 is the approximation of **X**, Σ_**X**_ is the (empirical) covariance matrix of **X** and 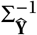 is the inverse of the (empirical) covariance matrix of the output of the original decoding model (see equation 1). When we solve for **W** in equation 2, we obtain:

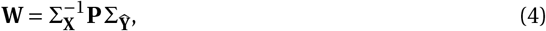

and observe that the weight matrix may be thought of as a combination of three subcomponents:

1. the covariance matrix of the data Σ_**X**_
2. the pattern matrix **P**
3. the normalizer 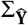

In the post-hoc framework, we replace the problem of finding the optimal weight matrix by the subproblems of finding the optimal Σ_**X**_, **P** and 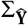. An initial estimate for the subcomponents can be obtained by applying a linear machine learning algorithm and decomposing its weight matrix using equation 2 (see also Figure 1). When understanding what the subcomponents represent and the subproblems they are trying to solve, the data analyst may use their domain information to refine the initial estimates at will. Afterwards, the modified subcomponents can be recomposed into an updated weight matrix (Figure 1):

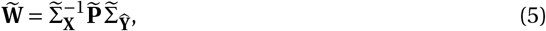

where 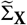 is a modified version of the data covariance, 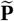 is a modified version of the pattern matrix, 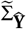 is a modified version of the normalizer, and 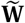 is the updated weight matrix that reflects the changes made to the subcomponents.

**Figure 1:**
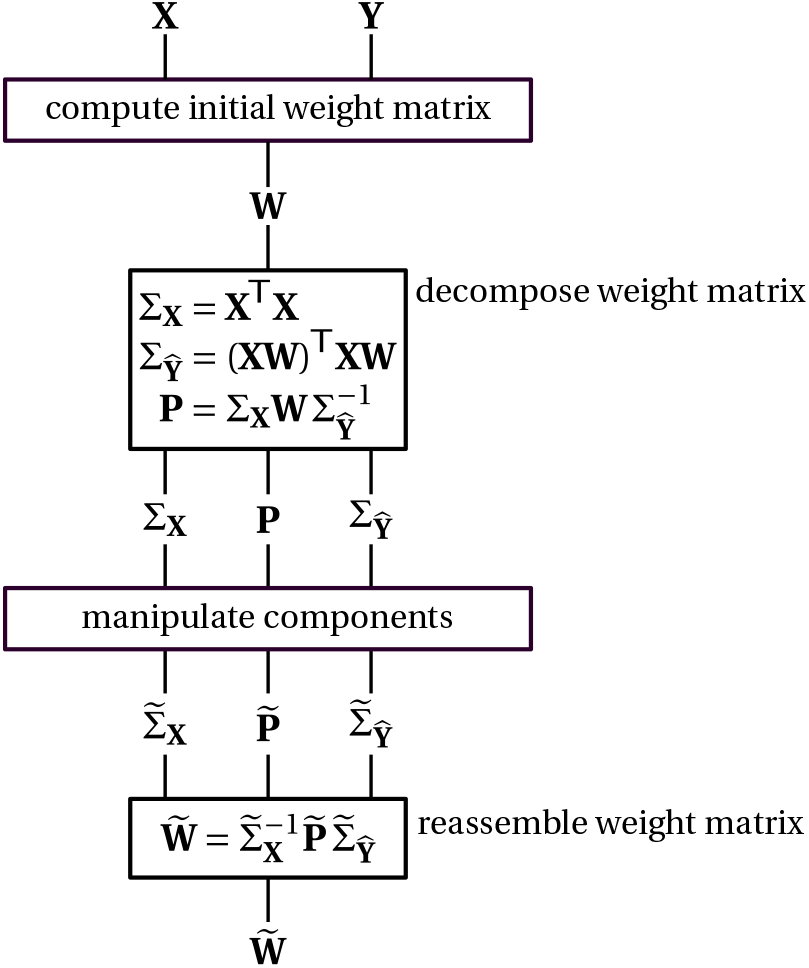
The post-hoc modification framework. First, the initial linear model **W** is constructed. This can for example be done with a general purpose linear machine learning algorithm. Then, using equation 2, **W** is decomposed into data covariance Σ_**X**_, pattern **P** and normalizer 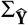. These subcomponents can then be manipulated at will to impose further restrictions on the model or inject prior information. Finally, the modified subcomponents 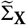, 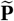 and 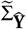 are reassembled into an updated linear model 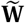.

We will now take a closer look at the three subcomponents. For a visual explaination, see Figure 2.

**Figure 2:**
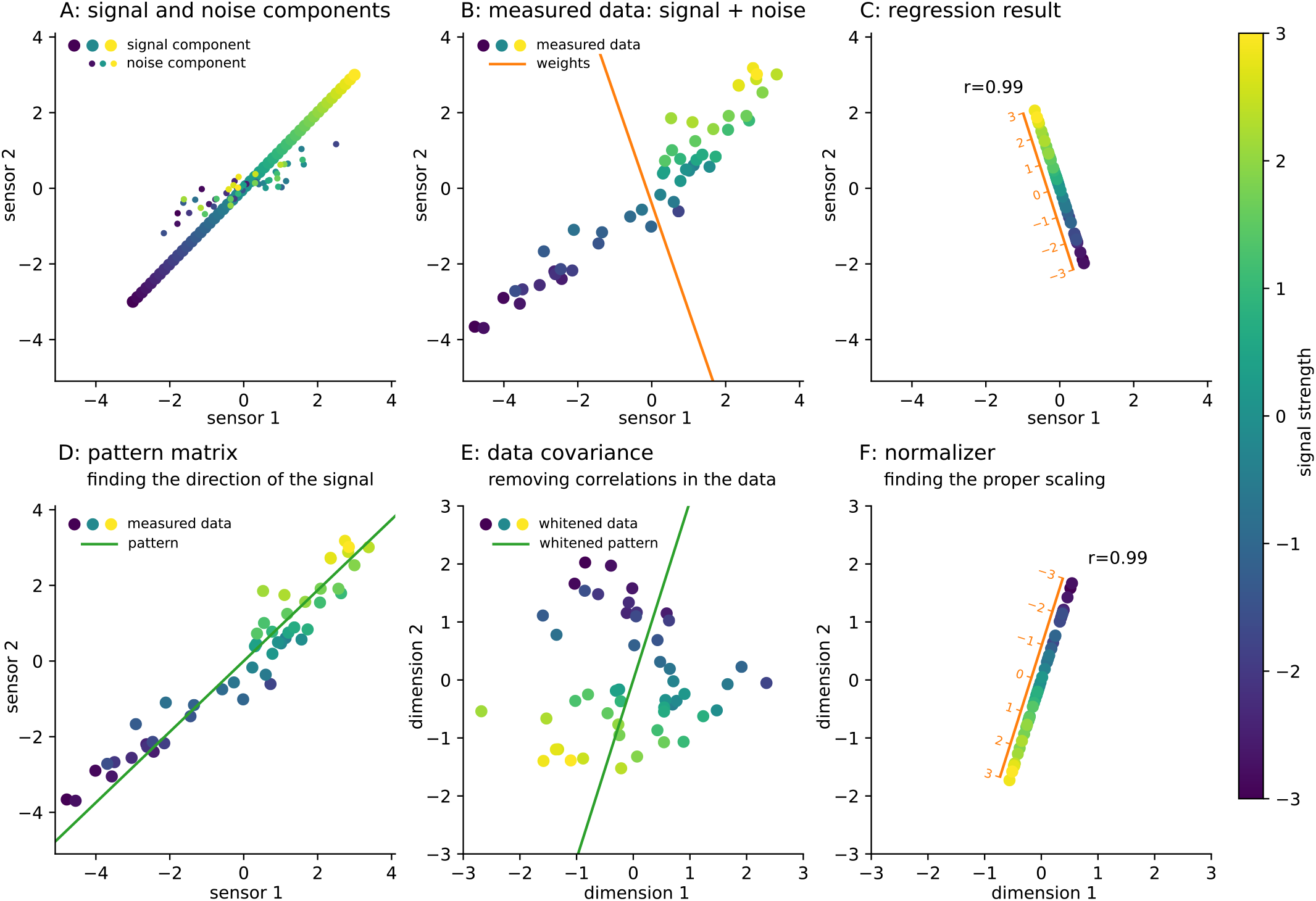
Visual explanation of the subcomponents of the post-hoc modification framework. This is a simulation of a signal that is being observed through two sensors. Dots represent observations of the signal and the color of the dots indicates the true signal strength during each observation. Linear regression is used to decode the true signal strength from the observed data. In visual terms, the task of the model is to decode the color of a dot, based on its location in the graph. **A:** The simulated data consists of two components. The first component (large dots) dictates how the signal is measured by the sensors (i.e. the encoding model). In this simulation, there is a one-to-one relationship between the true signal strength and the measurements at both sensors. The second component (small dots) is simulated using random numbers drawn from a two-dimensional Gaussian distribution and is a simulation of noise that is unrelated to the strength of the signal. **B:** The data that is recorded by the sensors (large dots) is the summation of both the signal and noise components. A linear regression model was trained on these observations, with the true signal strength as target, to determine the optimal linear transformation to map the measured data to signal strength. In this two-dimensional example, the model’s weights can be visualized as a line (orange line). We see that the direction of the regression line is dictated by the noise rather than the signal component, which is why the weight matrix is so hard to interpret. **C:** Applying the linear regression to the data is equivalent to projecting the measured data onto the regression line (orange axis). By projecting the data orthogonal to the noise, a near perfect reconstruction of the signal strength can be obtained. In the post-hoc modification framework, the model weights (orange line) are decomposed into three subcomponents, where each subcomponent solves a part of the regression problem. **D:** The pattern matrix represents the signal of interest and, like the weight matrix, can be visualized as a line (green). This line should approximate the direction of the actual signal (see panel A). **E:** The data covariance matrix is used to construct a whitening operator, which removes the correlations within the data. The data is projected such that all features are of unit variance and all cross-correlations between the features are eliminated. This transformation is then also applied to the pattern matrix (green line). Performing linear regression is now equivalent to projecting the whitened data onto the whitened pattern line. **F:** Finally, the normalizer (orange axis) scales the result such that the position along the projection line maps to the true signal strength. An interactive version of this figure is available at https://aaltoimaginglanguage.github.io/posthoc, where the noise component can be manipulated to study its effect on the subcomponents.

In order to design a mapping from **X** to **Y**, components of the data must be found that carry information that would be useful for determining the value of the decoding targets (Figure 2D, green line). Modifying the pattern matrix **P** allows for incorporating domain information on how the decoding targets **Y** are manifested in the data **X**.

To paraphrase de Cheveigné and Simon (2008), the filter needs to observe all components that “contaminate” the pattern components, so as to *subtract* them. Those observations may themselves be contaminated, requiring subtraction of additional components, and so on. The filter thus uses data from all input features, even the ones that carry no information about the decoding targets, in order to cancel out any contaminants in a delicate “balancing act” to get the best possible estimate of the pattern components. This is achieved by transforming the data such that all correlations between the input features are eliminated (Figure 2E), a process known as “whitening”. The 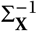 term in equation 4 represents a whitening transform that is applied to both the data and the pattern matrix (see appendix A). Modifying the data covariance matrix Σ_**X**_ allows for incorporating domain information on the correlations between the input features.

The “balancing act” described above attempts to eliminate any components that interfere with the pattern components. However, by itself, it does not untangle the pattern components from each other, nor impose a scaling on them. In the case of *k* = 1, the whitened data is projected onto the line that is defined by the whitened pattern matrix (Figure 2E, green line). In the case of *k* > 1, the pattern matrix defines a plane. As a final step, a mapping must be made between locations along the projection line/plane and the desired target **Y**. In the case of *k* = 1, this amounts to a scaling factor (Figure 2C, orange scale) and in the case of *k* > 1, the normalizer is a linear mapping between the locations on the projection plane to the model outputs **Y**. Modifying the normalizer 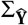 allows for fine-tuning of the relationship between the projected data and the decoding targets **Y**.

Domain information is by definition study specific, so in order to provide concrete examples, we will first introduce an example EEG study. In this study, the task of the linear model is to decode the forward association strength (FAS) between two words,^32^ based on an EEG recording of a participant reading the word-pair during a semantic priming experiment. We will then explore some ways in which the subcomponents may be modified to tune the model for this specific task.

### 2.3 EEG recordings

The decoding performance of two linear models was evaluated on an EEG dataset, which was recorded with 24 participants (7 female, aged 22–38, mixed handedness and all native speakers of Flemish-Dutch). Two recordings were dropped from the study: one was dropped due to problems with the stimulus synchronization signal and the other due to excessive sensor impedance. Participants signed an informed consent form prior to participating. Ethical approval of these studies was granted by an independent ethical committee (“Commissie voor Medische Ethiek” of the UZ Leuven, Belgium). These studies were conducted according to the most recent version of the declaration of Helsinki.

The participants read a series of sequentially presented words, organized in *prime*–*target* pairs, and pressed one of two mouse buttons to indicate whether the two words of a word-pair were related or not. The hand used to hold the mouse and the assignment of buttons to “yes”/“no” responses was counterbalanced across participants.

The prime word was presented for 200 ms and the target word for 2000 ms with a stimulus onset asynchrony (SOA) of 500 ms. Words were presented in white on a black background, rendered in the Arial font. Since a speeded button response task will generate ERP components that can mask N400 modulations,^33^ the participants were instructed to delay their button response until the target word turned yellow, which happened 1000 ms after the onset of the target word. The participants had 1000 ms to respond, or else a non-response code would be logged for the trial.

In addition to capturing the button response of the participant, EEG was recorded continuously using 32 active electrodes (extended 10–20 system) with a BioSemi Active II System, having a 5th order frequency filter with a pass band from 0.16 Hz to 100 Hz, and sampled at 256 Hz. An electro-oculogram (EOG) was recorded simultaneously using the recommended montage outlined by Croft and Barry (2000). Two final electrodes were placed on both mastoids and their average was used as a reference for the EEG.

### 2.4 Stimuli

The stimuli consisted of Flemish-Dutch word pairs (see section 2.11) with varying FAS between the two words in each pair, as measured by a large-scale norming study performed by De Deyne and Storms (2008). In this norm dataset, FAS is defined as the number of participants, out of 100, that wrote down the target word in response to the prime word in a free association task.

The stimuli used in the experiment were the top 100 word-pairs with highest FAS in the norm dataset and 100 word-pairs with an assumed FAS of zero that were matched in length, frequency and in-degree. Each word-pair with a high FAS consisted of words with a length of 3 to 10 letters, with no restrictions on frequency or in-degree. To construct the low FAS pairs, for each word in the high FAS condition, a random word was selected with equal length, frequency and in-degree (or, if no such word existed, a word that matched these as close as possible), and random pairings were made from the resulting words.

### 2.5 Data preprocessing

All data processing was performed using the MNE-Python^34^ and auto-reject^35^ software packages. The EEG was bandpass filtered offline between 0.1 Hz and 50 Hz by a 4th order zero-phase Butterworth filter to attenuate large drifts and irrelevant high frequency noise, but retain eye movement artifacts. Individual epochs were obtained by cutting the continuous signal from 0.2 s before the onset of each target stimulus to 1 s after. Baseline correction was performed using the average voltage in the interval before the stimulus onset (−200 ms to 0 ms) as baseline value. The random sample consensus (RANSAC) algorithm was used to detect excessively noisy channels, and those signals were subsequently replaced by interpolating the signals from nearby sensors using spherical splines.^36^ Two EOG artifact elimination passes were performed on the data. First, the EOG channels were used to attenuate eye artifacts from the EEG signal using the regression method outlined in Croft and Barry (2000). Second, the data was decomposed using independent component analysis (ICA) and any components that correlated strongly with one or more EOG channels were removed. Next, the signal was band pass filtered further using a tight passband around the frequency range in which the N400 component was found, namely between 0.5 Hz and 15 Hz, by a 4th order zero-phase Butterworth filter and downsampled to 50 Hz to reduce the dimensionality of the data. Further artifacts were removed using the autoreject procedure,^37^ which flags and interpolates noisy channels in each individual epoch by measuring how well data from other epochs predicts the data of the epoch currently under consideration. While autoreject can also flag and remove noisy epochs, this functionality was disabled to ensure no epochs were dropped from the data.

A full report of the data preprocessing steps can be found at: https://aaltoimaginglanguage.github.io/posthoc.

### 2.6 Initial linear models

In this paper, we give some examples on how to use the post-hoc modification framework to inject domain information into two general purpose machine learning models. For the regression scenario, we chose the ridge regressor as implemented in the Scikit-Learn package^38^ as the base model, and for the classification scenario the logistic regressor from the same package was chosen. These two particular models were chosen because they are widely used in neuroimaging and their performance on our example datasets is equal or better than other commonly used linear models (e.g. shrinkage linear discriminant analysis (LDA) or linear support vector machine (lSVM)).

Each epoch of the recording served as a single observation for the model, and the corresponding row-vector **x** was obtained by concatenating the timecourses recorded at all EEG sensors. The resulting vectors formed the rows of input matrix **X**, resulting in **X** ∈ ℝ^200×1600^. In the regression scenario, the desired output of the model, **Y** ∈ ℝ^200×1^, was specified as the log-transformed FAS of the word-pair presented during each epoch.^39^ In the classification scenario, **Y**was formed by specifying 1 if the word-pair presented during the epoch consisted of two associatively related words, and −1 otherwise.

Because we have a maximum of *n* = 200 epochs available for each participant, the problem of estimating 1600 weights from the data of a single participant is massively underspecified and the model will overfit.^40^ A common way to alleviate overfitting in linear models is to introduce regularization when estimating the covariance matrix during the training of the model. For example, with *ℓ*2 regularization, a trade-off is made between maximizing the fit between 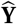 and **Y** and minimizing the absolute value of the weights ‖**W**‖, which prevents the model from placing too much emphasis on a single feature.^41 42^ Both initial models (ridge and logistic regression) implement such regularization. In the following subsections, we look at the problem of overfitting not from the perspective of the weight matrix, but from that of the subcomponents as defined by the post-hoc modification framework.

### 2.7 Strategies for modifying the covariance matrix

The data covariance matrix Σ_**X**_ is the subcomponent of a linear model that describes the (linear) relationships between the input features. Overfitting of the model will occurs when the linear relationships that were inferred from the training set do not hold on the test set, either because the estimation was incorrect or because the relationships change across observations (e.g. they change over time due to nonstationarity of the signal). In this case, the model will benefit from de-emphasizing the relations that were estimated on the train set in favor of a conservative ground truth that is expected to hold in both the training and test sets.

The *ℓ*2 regularization that is imposed on Σ_**X**_ by the initial models (ridge and logistic regression) adds a constant value to each diagonal element of the initial covariance matrix Σ_**X**_:

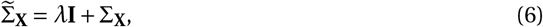

where **I** is an identity matrix of the appropriate size and *λ* ∈ [0 … ∞) is a parameter that controls the amount of regularization. One effect of this regularization scheme is that as *λ* approaches infinity, 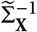 and hence 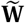 approach zero (equation 5). This effect is directly encoded in the optimization criterion for ridge regression.^43 44^ However, from the point of view of the subproblem that the covariance matrix represents, a second effect becomes apparent, namely that the covariance matrix is steered towards a scaled identity matrix. This means the model is steered towards a ground truth that none of the features are linearly related, meaning any of the relationships inferred from the training set are untrustworthy. It is this second effect that provides a straightforward insight into why *ℓ*2 regularization prevents overfitting and lends itself to schemes for incorporating domain information.

An approach that has the second effect, but not the first, is “shrinkage” regularization:^45 46^

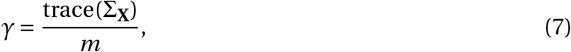

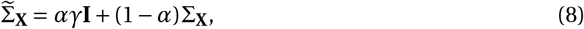

where *α* ∈ [0 … 1] controls the amount of shrinkage and *γ***I** is an identity matrix that is scaled by the mean of the diagonal elements of the empirical covariance matrix. In this regularization scheme, the covariance is steered towards a ground truth of no relationships between the features, without affecting the overall scaling of the matrix.

Both regularization schemes drive the covariance matrix towards a scaled identity matrix, penalizing all relationships equally in favor of the ground truth. One way of incorporating domain information is to distinguish between different kinds of relationships, and encode our belief that some may be estimated more reliably from the training data than others.

In our EEG example, **X** was obtained by concatenating the timecourses for each sensor. Such an approach to vectorizing the input data introduces a striking regularity in the covariance matrix, see Figure 3. The covariance matrix can be approximated by the Kronecker product^47^ between the spatial covariance matrix Σ_s_ (i.e., the linear relationship between the sensors) and temporal covariance matrix Σ_t_ (i.e., the linear relationship between the samples in time):^48^

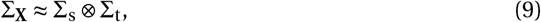

where (⊗) denotes the Kronecker product.

**Figure 3:**
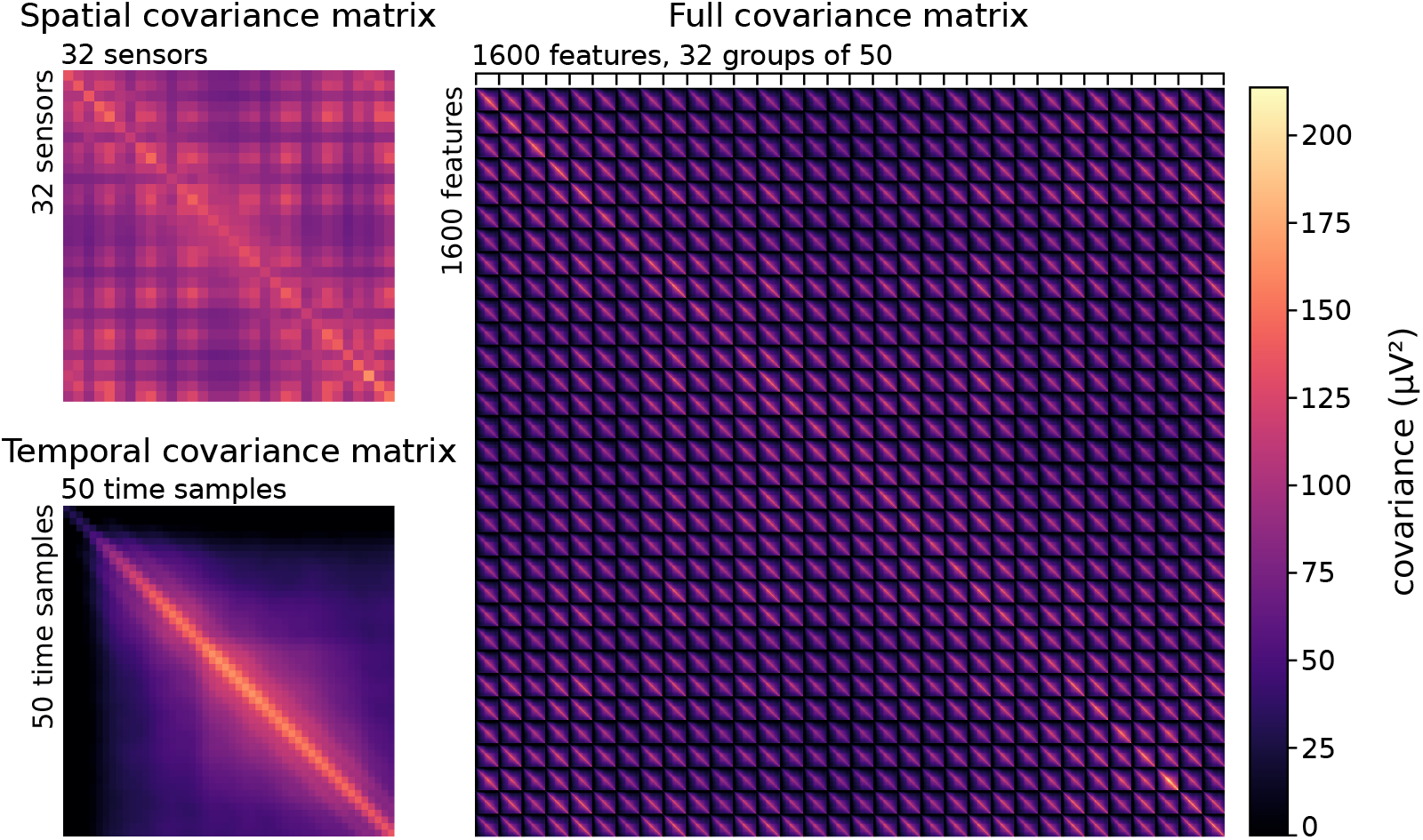
Shown on the right is the grand average covariance matrix. This matrix can be approximated with the Kronecker product of the grand average spatial covariance matrix (upper left) and grand average temporal covariance matrix (bottom left).

With this in mind, we propose a variation of the shrinkage approach that we call “Kronecker shrinkage”. First, we shrink of Σ_**X**_ towards Σ_s_ ⊗ **I**_t_, where **I**_t_ denotes an identity matrix of the same dimensionality as the temporal covariance matrix. Then, we substitute the result into equation 8 instead of Σ_**X**_:

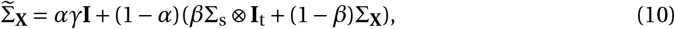

where *α* controls the shrinkage of the spatial component and *β* controls the shrinkage of the temporal component of the covariance matrix. This allows us to encode different amounts of confidence in the estimates of these two types of relationships from the training data.

### 2.8 Strategies for modifying the pattern matrix

The root problem that causes overfitting of the model is a lack of available training data. Therefore, for datasets that include multiple participants or recording sessions, one might expect that the model performs better if it had access to all recordings. However, in a neuroimaging setting, linear models that aim to generalize across participants are often outperformed by participant-specific models, even when the models have access to more training data.^49 50 51^ Since the optimal weights depend on both the signal of interest and any interfering signals, it is often not straightforward to transfer a weight matrix from one participant to another.

The pattern matrix Σ_**P**_ is the subcomponent of a linear model that describes only the signal components that are informative of the targets, as opposed to other “noise” components. In some cases the pattern matrix is likely to be similar across participants. In our example study, the task was to decode FAS from the EEG signal, in which case the literature notes the N400 component of the ERP ^52 53^ as the primary signal of interest. While there are factors that affect the latency of this component, such as age,^54^ the participants in our example study were drawn from a homogeneous pool (university students), so we can expect the timing of the component, as well as its distribution across sensors, to be relatively stable. Also in the case of other, similar N400 studies, the pattern matrix has been successfully transferred between participants.^55 56^ Hence, a good strategy for improving the estimation of the pattern matrix may be to bias it towards a grand-average pattern matrix that was obtained from the recordings of other participants.

Let 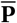 be the average of the pattern matrices for all recordings, excluding the recording currently under consideration. Then:

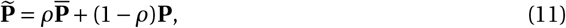

where *ρ* controls how much the pattern matrix is steered towards the grand average. This operation can be beneficial if the model has difficulty identifying the signal of interest during the training phase (e.g, due to noisy data, lack of training data, or absence of a **Y** matrix^57^).

Another approach to correcting inaccuracies in the pattern matrix is to leverage the fact that in our semantic priming study, the signal of interest (the N400) is well localized in time. One way of achieving this would be to restrict the data **X** to a time window surrounding 400 ms. However, this would deprive the model from potentially useful observations of the noise components that the model is attempting to cancel out. A good example can be found in the domain of EEG/MEG source estimation, where, even if the goal is to estimate activity at a single dipole source, it is beneficial to create a spatial filter using many sensors, and not only the sensors that are most sensitive to activity at the source dipole.^58^ The post-hoc modification framework allows us to place restrictions on the pattern timecourses alone, keeping information about the noise components intact.

In our example study, we multiplied the timecourses in the pattern matrix with a Gaussian kernel (Figure 4):

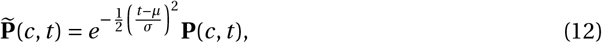

where *c* iterates over all channels, *t* iterates over all time samples, and **P**(*c*, *t*) denotes the element of **P** that corresponds to channel *c* at time *t*. Parameters *μ* and *σ* determine the center and width of the Gaussian kernel (Figure 4).

**Figure 4:**
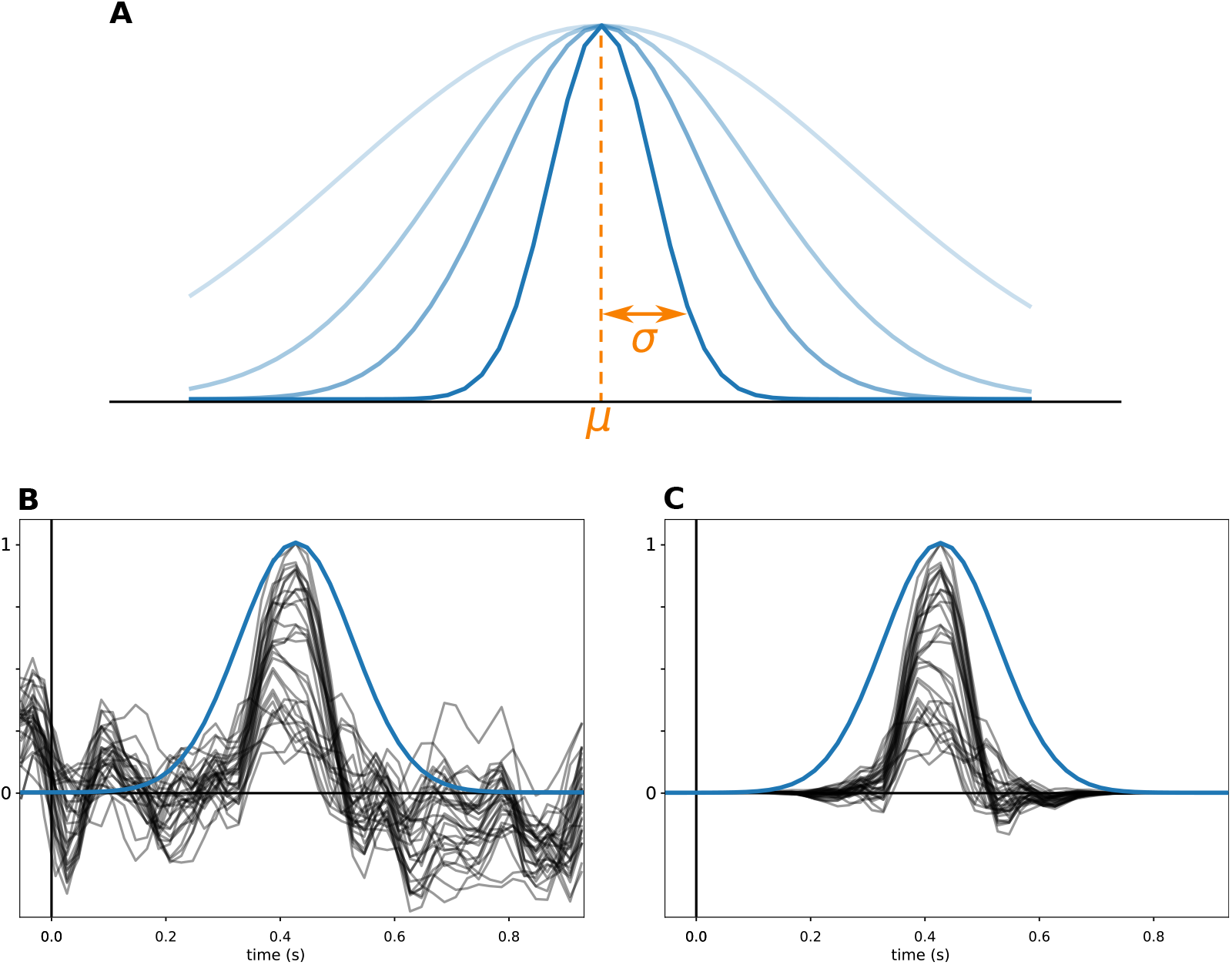
Example of multiplying the pattern matrix with a Gaussian kernel. **A:** Parameters *μ* and *σ* determine the position and shape of the kernel. **B:** Example of a pattern matrix, with the timecourse for each sensor drawn in black. An example Gaussian kernel is drawn in blue. For this visualization, the pattern was normalized to have a maximum amplitude of 1 to have the same visual scale as for the kernel. **C:** The result of multiplying the pattern matrix with the Gaussian kernel.

### 2.9 Strategies for modifying the normalizer

Modifications to the covariance and pattern matrices result in changes to the projection line (*k* = 1) or plane (*k* > 1) of the model. This means that the normalizer needs to be recomputed to re-map locations along the projection line/plane to the model outputs.

One way to compute an appropriate normalizer is to find the least-squares mapping between the output of the “raw” filter 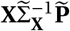 and **Y**, through linear regression:

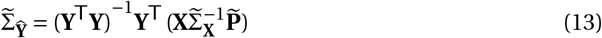

### 2.10 Model evaluation and automated tuning of the hyperparameters

The performance of each model was evaluated for each participant separately, using 10-fold crossvalidation. The order of the observations in the recording (the rows of **X** and **Y**) were shuffled and then assigned to ten folds. Two crossvalidation loops were used, which we will refer to as the “outer” and “inner” loops.

In the outer crossvalidation loop, nine folds were used as training data and one fold was used as test data. Normalization of **X** was performed inside the outer crossvalidation loop, such that the mean and standard deviation of each feature across observations was computed on the training data only, and subsequently used to normalize the features of the test data. By repeating this ten times, such that each fold has been used as test data once, and collecting the output of the model for each run, the full matrix 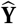 was constructed, containing the crossvalidated model output for each epoch. The performance of the model, *p*, was then quantified in the regression scenario using the Pearson correlation between 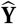 and **Y**, and in the classification scenario using the classification accuracy.

When a model incorporates data from other recordings (the “multiple subjects” and “all information” models, see section 2.8), a distinction was made between the recording for which the model was currently being evaluated and the recordings made on the other participants. During the outer crossvalidation loop, the training data was augmented with the data from the other participants, while the test data was left untouched.

Both initial models (see section 2.6) have a parameter (*α*) that determines the amount of *ℓ*_2_ regularization, and throughout sections 2.7 to 2.8, we have defined several more parameters (*β*, *ρ*, *μ*, *σ*) that control various aspects of the model. These parameters can be used to impose hard constraints on the model, for example, *μ* and *σ* limit the time-range in which the model will search for the signal of interest. Alternatively, they can be treated as parameters that need to be learned, just like the model weights.

In our example analysis, we used an “inner” leave-one-out cross-validation loop to learn these parameters during the training phase. Since searching the entire parameter space would be too time consuming, we first evaluated 100 random values for the parameters, taking the best performing parameter set as rough first estimate. This estimate was then fine-tuned using a convex optimization algorithm (Limited-memory Broyden–Fletcher–Goldfarb–Shanno with box constraints (L-BFGS-B)^59^). This algorithm searches for the optimal parameters by alternating between two phases: 1) estimating the direction of maximum performance gain by making tiny changes to each parameter and measuring the effect on the leave-one-out performance of the model, followed by 2) updating the parameters in the direction of maximum positive effect on the performance. This process is repeated until no changes to the parameters can be found that improve the leave-one-out performance.

The optimization approach employed by the L-BFGS-B algorithm requires that the chosen model performance evaluation function is continuous and differentiable. This is why, for the classification model, we used the logistic loss function rather than classification accuracy or receiver operating characteristic – area under curve (ROC-AUC), since the latter two are not differentiable. For the regression model, Pearson correlation between the leave-one-out model output and the desired output (**Y**) was used as a loss function, as this is the measure we report in the results section. This measure is closely related to the more traditional mean squared error (MSE) loss function, but is easier to interpret, as it has been normalized to range from 0 to 1.

### 2.11 Data and code availability

Electronic supplementary information is available at: https://aaltoimaginglanguage.github.io/posthoc. This includes a Python package that provides an implementation of the post-hoc modification framework that is compatible with Scikit-Learn.^60^ The package contains optimized implementations (see appendices B and C) of all modification strategies discussed in this paper and provides an interface for implementing new ones.

The consent form that was signed by the participants stated that the raw data would not be shared publicly without limitations. This data can be obtained upon request from the corresponding author, for reasons such as replication, assessment of other analysis methods, or aid in future studies on semantic processing.

All nonsensitive data can be found in the electronic supplementary information, including the grand-average pattern matrices, the preprocessing reports for the data of each participant, the output of the models and the stimulus list.

## 3 Results

We determined the effectiveness of the strategies for incorporating domain information by comparing the performance of the models that incorporates domain information to that of the original models. See Table 2 for an overview of the models that were evaluated.

**Table 2:**
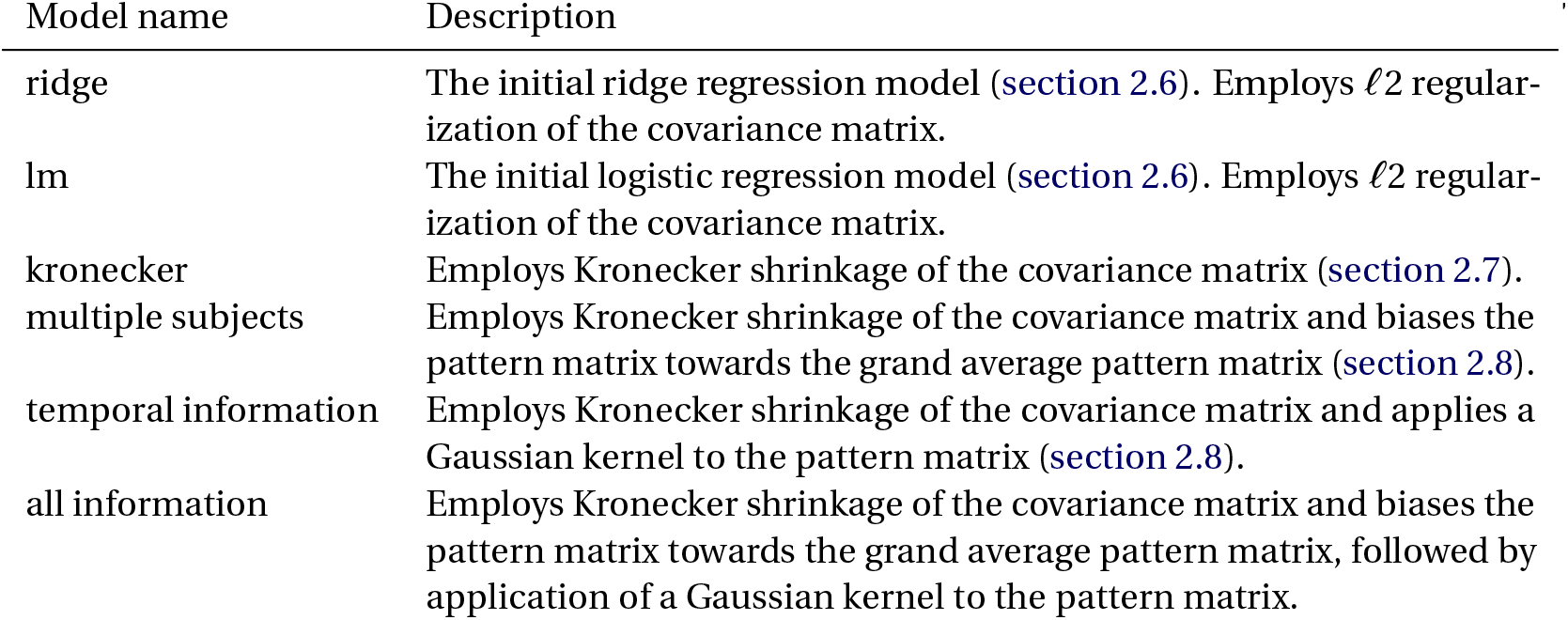
Models that were evaluated

The performance of the models was evaluated using 10-fold cross validation (the epochs were shuffled before being assigned to folds) and presented in Figure 5. For regression models, we report the Pearson correlation between the model output and the FAS of the word-pairs as the performance metric (Figure 5A). For classification models, we report the classification performance using the ROC-AUC score (Figure 5B), where the classification task was to assign each epoch to either the low-FAS or high-FAS category.

**Figure 5:**
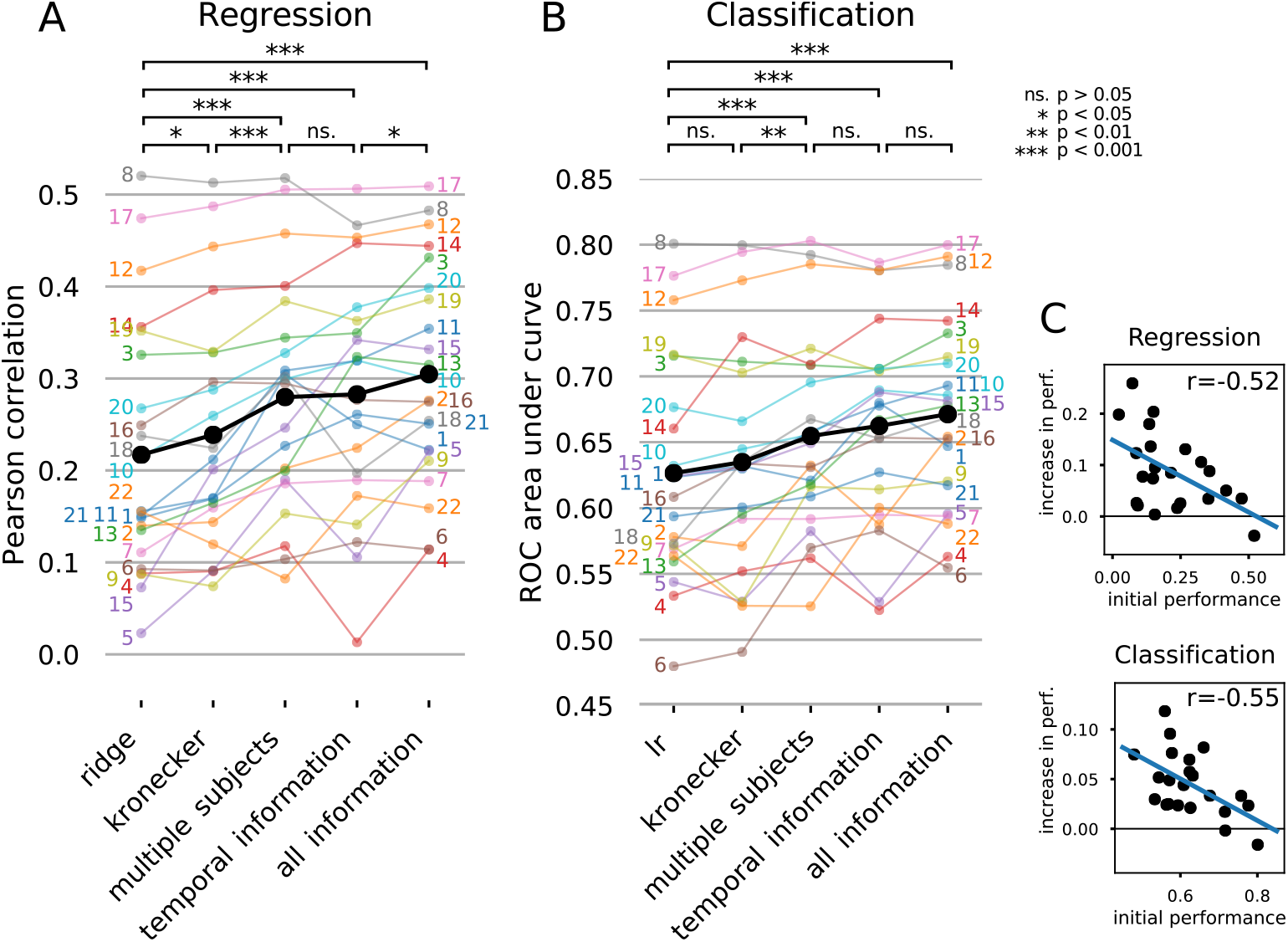
Performance of the linear models, before and after applying various post-hoc modification strategies. The performance for each participant is shown (colored dots and numbers), along with the mean performance across participants (black dots). Lines have been drawn between the dots in order to facilitate comparing the performance of a single participant across modification strategies. At the top, statistical comparisons between the group-level performances of the methods (paired t-tests) are shown. See the main text for an explanation of the modification strategies. **A**: Performance of the regression model. **B**: Performance of the classification model. **C**: The relationship between the performance of the initial model and the increase in performance gained by including domain information (the “all information” model).

Taken individually, each manipulation strategy provided a small improvement to the performance of the initial model (for statistics, see top of Figure 5). Taken together (the “all information” model), the performance was substantially improved by using post-hoc modification to inject domain information for both the initial ridge regression (effect size: 0.088, pair-wise *t*-test: *t* = 5.526, *p* < 0.001) and logistic model (effect size: 0.045, *t* = 6.550, *p* < 0.001).

The post-hoc modification strategies for incorporating domain information were set up such that the model could always fall back to not incorporating any domain information. Hence, in theory, the models should incorporate domain information only when it is beneficial. Inspecting the optimized parameters (Table 3) reveals which types of domain information were incorporated by the model. In practice, the models optimized their parameters based on the training set only, using an inner cross-validation loop, hence can be suboptimal for the test set due to overfitting. Indeed, for participant 8, where the initial models performed best, incorporating domain information proved detrimental (Figure 5, gray lines). Generally, for recordings on which the initial models had low performance, the models had the most to gain from incorporating domain information, with diminishing returns for cases in which the initial model was already performing well (Figure 5C).

**Table 3:** Optimal parameters for the “all information” model

| subject | Regression |  |  |  |  | Classification |  |  |  |  |
| --- | --- | --- | --- | --- | --- | --- | --- | --- | --- | --- |
| | $\alpha$ | $\beta$ | $\rho$ | $\mu$ (s) | $\sigma$ (s) | $\alpha$ | $\beta$ | $\rho$ | $\mu$ (s) | $\sigma$ (s) |
| 1 | 0.97 | 0.11 | 1.00 | 0.77 | 0.96 | 0.60 | 0.07 | 0.50 | 0.42 | 0.24 |
| 2 | 0.71 | 0.01 | 0.21 | 0.38 | 0.08 | 0.74 | 0.00 | 0.28 | 0.41 | 0.24 |
| 3 | 0.90 | 0.03 | 0.15 | 0.37 | 0.07 | 0.66 | 0.02 | 0.24 | 0.37 | 0.07 |
| 4 | 0.94 | 0.36 | 0.32 | 0.40 | 0.32 | 0.97 | 0.72 | 0.52 | 0.75 | 0.90 |
| 5 | 0.70 | 0.00 | 0.64 | 0.41 | 0.24 | 0.80 | 0.00 | 0.65 | 0.41 | 0.24 |
| 6 | 0.03 | 0.03 | 0.26 | 0.65 | 0.38 | 0.01 | 0.07 | 0.99 | 0.10 | 0.35 |
| 7 | 1.00 | 0.35 | 0.22 | 0.30 | 0.78 | 0.99 | 0.00 | 0.00 | 0.30 | 0.78 |
| 8 | 0.54 | 0.00 | 0.11 | 0.47 | 0.94 | 0.46 | 0.00 | 0.12 | 0.47 | 0.94 |
| 9 | 0.82 | 0.00 | 0.23 | 0.38 | 0.07 | 0.92 | 0.00 | 0.62 | 0.22 | 0.95 |
| 10 | 0.96 | 0.02 | 0.82 | 0.22 | 0.95 | 0.80 | 0.01 | 0.32 | 0.41 | 0.24 |
| 11 | 0.24 | 0.02 | 0.36 | 0.66 | 0.37 | 0.22 | 0.02 | 0.19 | 0.66 | 0.37 |
| 12 | 0.61 | 0.01 | 0.22 | 0.66 | 0.71 | 0.26 | 0.00 | 0.12 | 0.66 | 0.71 |
| 13 | 0.64 | 0.00 | 0.31 | 0.37 | 0.07 | 0.64 | 0.00 | 0.23 | 0.37 | 0.07 |
| 14 | 0.05 | 0.18 | 0.14 | 0.38 | 0.10 | 0.02 | 0.04 | 0.08 | 0.08 | 0.69 |
| 15 | 0.86 | 0.08 | 0.00 | 0.38 | 0.08 | 0.75 | 0.05 | 0.03 | 0.38 | 0.09 |
| 16 | 0.18 | 0.01 | 0.03 | 0.66 | 0.71 | 0.17 | 0.01 | 0.02 | 0.66 | 0.71 |
| 17 | 0.53 | 0.04 | 0.29 | 0.24 | 0.39 | 0.32 | 0.01 | 0.27 | 0.08 | 0.69 |
| 18 | 0.43 | 0.13 | 0.36 | 0.42 | 0.44 | 0.50 | 0.03 | 0.38 | 0.74 | 0.88 |
| 19 | 0.48 | 0.02 | 0.31 | 0.47 | 0.94 | 0.30 | 0.01 | 0.19 | 0.47 | 0.94 |
| 20 | 0.63 | 0.05 | 0.81 | 0.29 | 0.10 | 0.58 | 0.03 | 0.38 | 0.38 | 0.08 |
| 21 | 0.90 | 0.17 | 0.61 | 0.00 | 0.82 | 0.85 | 0.06 | 0.46 | 0.40 | 0.31 |
| 22 | 0.75 | 1.00 | 0.41 | 0.64 | 0.05 | 0.55 | 0.42 | 0.29 | 0.73 | 0.26 |

We will now look more closely into the effectiveness of the individual strategies.

One factor that influences the performance of the model is the amount of noise and the ability of the model to accurately determine the “direction” of the noise (see Figure 2). By applying regularization to the covariance matrix, the estimated direction of the noise is steered towards being spherical (i.e. equal in all directions). Both initial models already apply *ℓ*_2_ regularization. In the regression scenario, Kronecker shrinkage (Figure 5, left, “kronecker”), which controls the amount of shrinkage for the spatial and time dimensions separately, outperforms the *ℓ*_2_ regularization approach (paired *t*-test: *t* = 2.81, *p* < 0.05). In the classification scenario, Kronecker shrinkage is beneficial in some cases, but detrimental in others (Figure 5, right, “kronecker”) and does not significantly outperform *ℓ*_2_ regularization (*t* = 1.48, *p* > 0.05).

Table 3 lists the parameters chosen by the “all information” model, for each subject, fitted to the entire dataset. Heavy shrinkage is applied by most models (high values for *α*), however, many models made little use of shrinking the temporal component of the covariance matrix (low values for *β*).

Inspecting the pattern matrices (Figure 6), computed with equation 2, reveals another contributing factor that influences the performance of the models. The N400 component is a prominent signal of interest for determining FAS from EEG data.^61^ In some patterns (e.g, participants 3 and 20), the N400 is clearly visible as a peak at around 400 ms. However, in almost all patterns, there are other peaks, indicating that the model has learned other signals of interest as well. The question is how well these features generalize beyond the training set.

**Figure 6:**
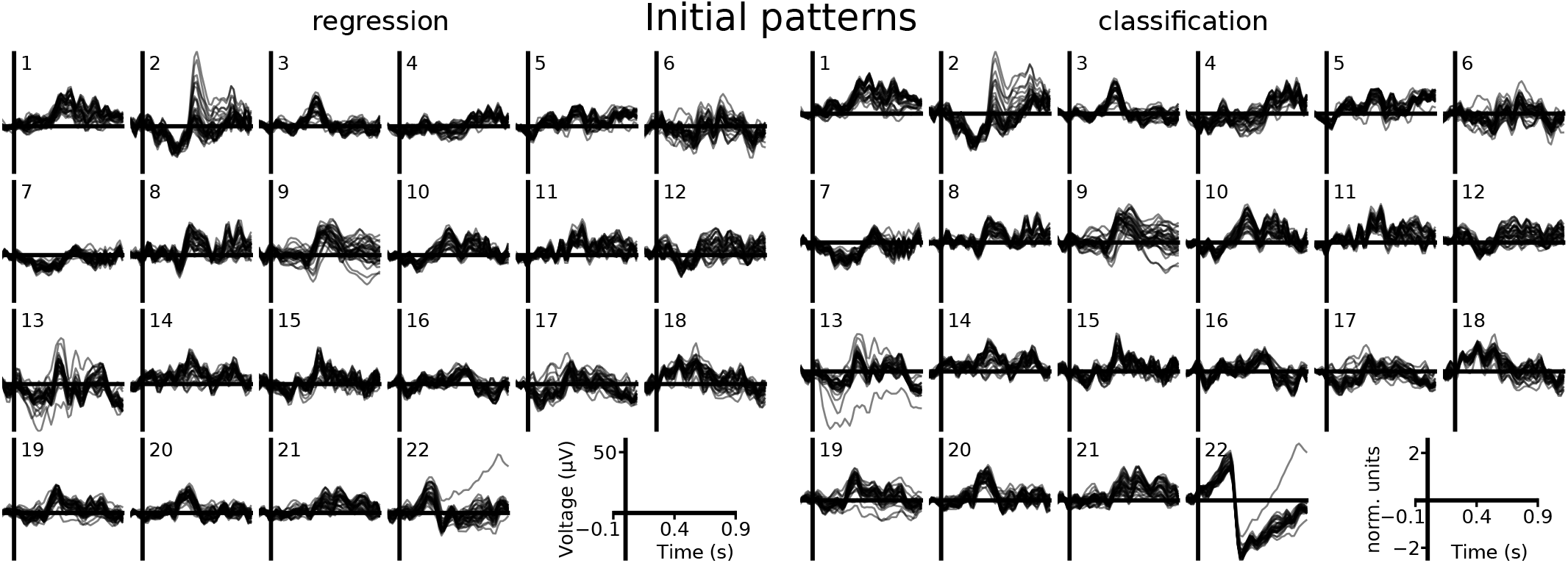
For each participant (1-22), the pattern that was learned by the initial linear models, for both the regression (left, ridge regression) and classification (right, logistic regression) scenarios. The timecourses of all electrodes are shown overlaid.

We introduced two strategies to bias the pattern matrix towards isolating the N400 component. First, a template of the N400 component was constructed by computing the grand-average pattern across participants other than the one currently being analyzed. Taken in isolation, the “multiple subjects” strategy improved the model beyond the “kronecker” model, both in the regression (paired *t*-test: *t* = 4.89, *p* < 0.001) and classification (*t* = 3.27, *p* < 0.01) scenarios. Second, the pattern was limited in time, allowing the model to focus on a single ERP component. Taken in isolation, the “temporal information” strategy performed equally well, both in the regression (vs. “kronecker”: *t* = 3.58, *p* < 0.01 vs. “multiple subjects”: *t* = 0.24, *p* = 0.81) and classification (vs. “kronecker”: *t* = 3.77, *p* < 0.01 vs. “multiple subjects”: *t* = 1.10, *p* = 0.29) scenarios. When both strategies were applied in tandem (“all information”), performance was increased even further, compared to the “multiple subjects” model, in both the regression (*t* = 2.39, *p* < 0.05) and classification (*t* = 3.81, *p* < 0.01) scenarios. Compared to the “temporal information” model, the “all information” model’s performance was significantly better than the “temporal information” model only in the regression scenario (*t* = 2.54, *p* < 0.05).

Looking at the “all information” model, for most participants, the optimizer chose to bias the pattern matrix towards the grand-average (Table 3, high values for *ρ*). Then, for a selection of participants, the optimizer chose to further refine the pattern by restricting it to a narrow time window surrounding the N400 component (Table 3, *μ* around 400 ms and low values for *σ*). Overall, the optimized patterns show a much more pronounced N400 effect (Figure 7) compared to the patterns of the initial models (Figure 6), indicating that the N400 was indeed a stable feature of interest that generalizes well beyond the training set. For some participants, the initial models failed to find a signal that clearly resembles the N400 potential, yet when a template N400 signal was mixed in with the pattern matrix, the decoding accuracy increased, which suggests that the N400 potential was present in the EEG of the participant after all (e.g., compare Figure 6 and Figure 7 for participants 5 and 13).

**Figure 7:**
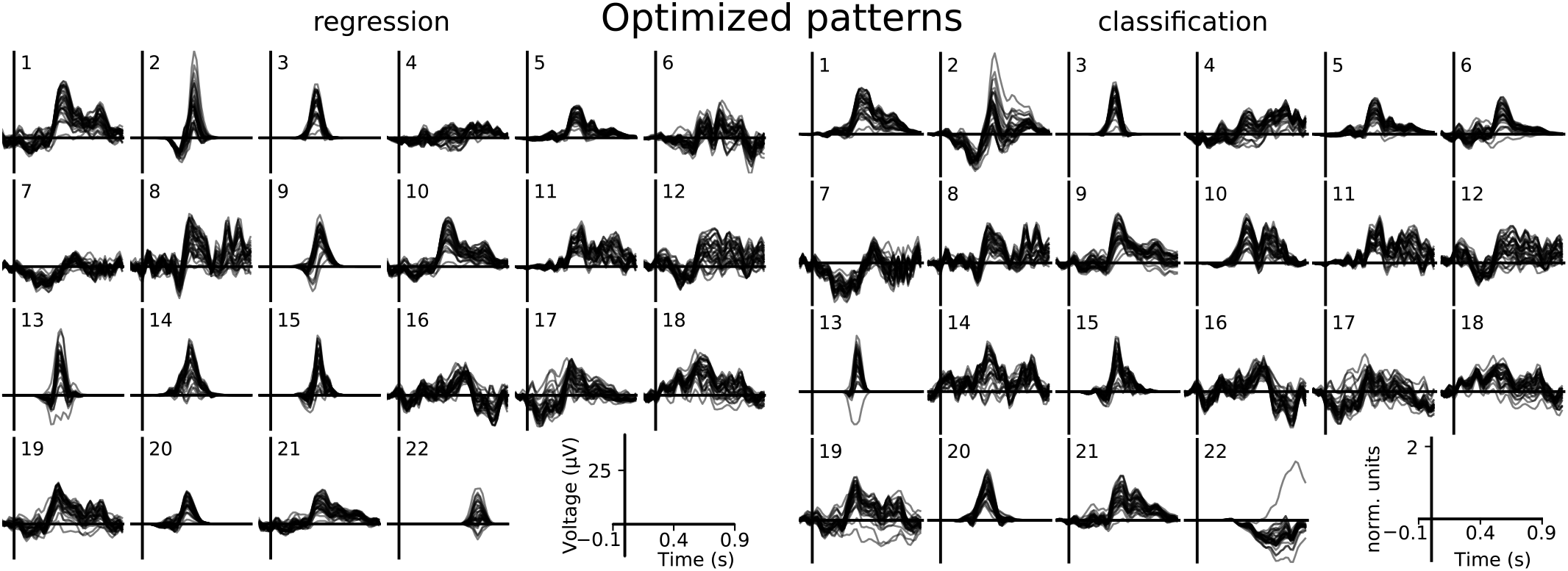
For each participant (1-22), the pattern that was used in the linear model that incorporates all post-hoc modifications (the “all information” model), for both the regression and classification scenarios. The timecourses for all electrodes are shown overlaid.

## 4 Discussion

We have demonstrated how domain information can be incorporated into general purpose linear models with the post-hoc modification framework. When using this framework, we shift our focus away from estimating a weight matrix towards the subproblems of 1) modeling the signal of interest (the pattern matrix), 2) establishing the relationship between input features (the data covariance) and 3) performing a normalization step.

As Haufe et al. (2014) pointed out, there is a strong parallel between the pattern matrix and the concept of a leadfield or “forward solution”, as used in source estimation.^62^ From this perspective, the decoding targets are similar to the source dipoles and the weight matrix is similar to the inverse operator. The main difference is that the pattern matrix is not constructed by modelling volume conduction in the head, but rather through a linear machine learning algorithm. In this work, we have extended the parallel further by observing that the domain of source estimation has always approached the computation of the inverse operator (or spatial filters) as a multi-step process, where first the covariance matrix is computed on the sensor data, which is then combined with the leadfield,^63^ and we may use the same approach when fitting decoding models.

From this point of view, possibilities for incorporating domain information into the model become obvious. In this work, we have explored a few possibilities to modify a ridge regression and logistic regression model to:

1. employ Kronecker shrinkage that takes the spatio-temporal nature of EEG into account
2. use the grand-average pattern across multiple recordings as a prior for the current model
3. use information about the temporal characteristics of the N400 potential as a further prior

The resulting models show a remarkable improvement over the initial general purpose models (Figure 5).

The post-hoc modification framework opens up a wide range of possibilities to design strategies for incorporating domain information. Our examples aim to demonstrate the capabilities of the framework and serve as inspiration for designing new strategies for other study paradigms or recording modalities.

One may explore more informative priors for the covariance matrix than an identity matrix. For example, bandpass filtering the signal will introduce a predictable dependency between consecutive time samples, which may be used as a shrinkage target for the temporal component of the covariance matrix. Likewise, for EEG and MEG studies, volume conduction in the head will impose a predictable dependency between the signals at different sensors, which can be modeled using a leadfield.^64^ Also for the pattern matrix, there are other avenues of domain information to explore. For example, the N400 potential has a well defined spatial signature^65^ that may be used as a prior for the pattern matrix. Finally, there might also be opportunities to incorporate domain information through the normalizer, although we did not explore this in this study and treated the normalizer as a mere scaling of the model output. Inspiration for normalization schemes can be found in the beamformer literature.^66^ For example, if the pattern matrix has been crafted to be in some measurement unit, one may wish to enforce that model output adheres to the same unit. The unit-gain constraint the of the linearly constrained minimum variance (LCMV) beamformer, **WP** = **I**, ensures that units are preserved. Using post-hoc modification, we can apply the unit-gain constraint of the LCMV beamformer to any linear model by using:

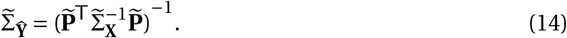

A common approach to reducing data dimensionality is to first apply a spatial filter, followed by a temporal filter.^67 68 69^ While the resulting model becomes blind to interactions happening in a different locations at different times, the reduction in dimensionality will decrease overfitting, potentially offsetting the disadvantages. Such an approach can be explored in the post-hoc framework as well, with the benefits that the choice of whether to treat space and time separately or jointly no longer has to be made model-wide, but can be done for each subcomponent separately. For example, the empirical covariance matrix can be replaced with the Kronecker product of the spatial and temporal covariance matrices, and the pattern matrix can be replaced with the outer product of a spatial and temporal pattern, for example obtained using non-negative matrix factorization.^70^ As in our example modifications, a hyperparameter can be defined to scale the matrices between the full spatio-temporal forms and the reduced forms that treat space and time separately, allowing the model to dynamically seek out the most suitable approach.

In our examples, we optimized the hyperparameters (*α*, *β*, *ρ*, *μ*, *σ*) using only the decoding performance of the resulting model as performance metric, but one can imagine using other metrics. For example, decoding models are often employed to explore the signal of interest that was learned, in which case interpretability of the model is more important.^71 72^ In this case, one may wish to optimize a tradeoff between sparsity of the pattern matrix (not to be confused with sparsity of the weight matrix) and decoding performance.^73^

Furthermore, the fact that a signal is useful for a decoding task does not necessarily mean that it is of interest to the study. For instance, in our example EEG study, eye artifacts can be a predictor for FAS ^74 75^ and, despite the preprocessing steps to attenuate them, are likely still present in the pattern matrices (e.g., Figure 6, participant 22). Furthermore, given that most models in neuroimaging are overfitting due to the ratio of number of features versus the size of the training set, the pattern matrix can be noisy and/or biased.

If the goal of the analysis is to study a specific signal of interest, it may be desirable to fix aspects of the pattern. For example, if the goal is to measure the timing of the N400 potential, we may explicitly set the pattern matrix to a time-shifted version of a suitable N400 template.

Restricting the pattern allows for precise control over which aspects are “learned” from the data and which are dictated by the researcher. If 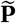 is completely fixed, the model is transformed into a beamformer^76^ and no ground truth (**Y**) is required to train the model. For example, it is possible to train a model on a dataset for which a ground truth is available, and transplant the resulting pattern matrix into a new model that is fitted to a dataset for which no ground truth is (yet) available.^77^

Taking the opposite view, one may wish to use the post-hoc modification framework to steer the model away from signals that are known to be relevant for the decoding task, in order to force the model to explore as yet unknown signals. In this case, the known signals of interest may be removed from 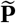, which will result in this signal being explicitly tagged as noise to be filtered out.

While the above examples are all in the domain of machine learning, linear models are also widely used in the domain of statistics, where applications range from familiar t-tests, through ANOVA F-tests, to more advanced multilevel models. The post-hoc framework can by applied here as well. For example, the “multiple subjects” model, which biases the pattern matrix to a group average, parallels a linear mixed-effects model which performs a similar trick to compute both a group-level slope as well as slopes for individuals.^78^

We envision the post-hoc modification framework as an iterative process, where an initial model is fitted to the data without any restrictions. This is followed by an inspection of the resulting patterns, covariance and normalizer by the data analyst, who then proceeds to place restrictions using post-hoc modification. The model is fitted again, taken the new restrictions into account and the cycle continues until finally, a model is obtained that satisfies all requirements of the study. In this manner, machine learning becomes less of a “black box” and more a dialogue between data analyst and model.

## 5 Conclusion

In the post-hoc modification framework, the weight matrix of a linear model is regarded as a combination of three subcomponents: a pattern matrix, a data covariance matrix, and a normalizer. The problem of computing a weight matrix can accordingly be split up into the subproblems of estimating each subcomponent. We showed how domain information can often be straightforwardly formulated in terms of these subcomponents. An initial estimate for the subcomponents can be obtained by decomposing the weight matrix as produced by a linear machine learning algorithm. In what we call “post-hoc modification” each subcomponent can then be refined at will, which provides opportunities to incorporate domain information. Afterwards, the modified subcomponents are re-assembled into a weight matrix, which now incorporates the injected domain information.

We have presented some strategies for incorporating domain information and demonstrated their effectiveness on an example EEG dataset, where the task of the linear model was to predict, given a single epoch, the associated relatedness between the two words that were presented during the epoch. Through post-hoc modification of two general purpose models, a ridge regression and logistic regression model, information was incorporated about the spatio-temporal nature of EEG data, the recordings performed on other participants, and the N400 potential. The resulting domain specific models achieved an increase in decoding performance compared to the initial, general purpose models.

However, as domain information is study specific, so are post-hoc modification strategies. While some of the presented strategies can be appropriate for other EEG studies, they mainly serve as examples of how the post-hoc modification framework offers many possibilities to implement modification strategies to suit the many different purposes of linear models in neuroimaging and other fields.

## 6 Acknowledgements

The EEG data was recoded at the KU Leuven, Department of Neurosciences, under the supervision of Marc Van Hulle.

MvV was supported by the Interuniversity Attraction Poles Programme – Belgian Science Policy (IUAP P7/11) and a grant from the Aalto Brain Centre (ABC), and is currently supported by the Academy of Finland (grant 310988). RS is supported by the Academy of Finland (grant 315553) and the Sigrid Jusélius Foundation.

## Appendix A The relationship between 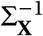 and whitening

The 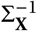 term in equation 4 represents a whitening transform that is computed using **X** and subsequently applied to both the data **X** and the pattern matrix **P**. This becomes clear when we rewrite 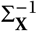 in terms of the eigendecomposition of Σ_**X**_:

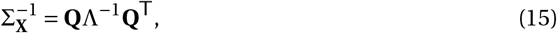

where **Q** is a matrix where each row is an eigenvector of Σ_**X**_ and Λ is a diagonal matrix where each diagonal element is the corresponding eigenvalue. Then, the linear transformation Φ that whitens **X** is defined as:

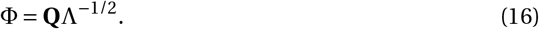

Hence, 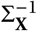 can be rewritten as:

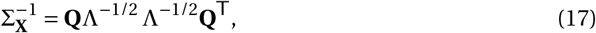

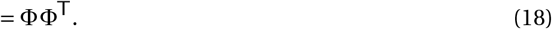

and we can show that that when the model is applied, it performs a whitening transformation on both the data **X** and the pattern matrix **P**:

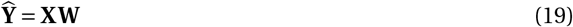

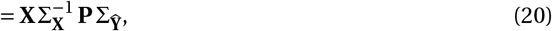

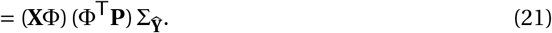

## Appendix B Optimizing covariance computation

Computing the empirical covariance matrix Σ_**X**_ and its inverse 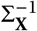 can be time consuming, given the number of features in EEG and especially MEG epochs. Typically, however, the number of features far exceeds the number of epochs, which allows us to compute equation 5 efficiently by applying the matrix inversion lemma,^79^ which states that for any matrices **A**, **B**, **U**, and **V** of appropriate size, the following holds:

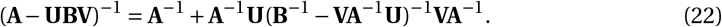

This allows us to reformulate **X**^T^**X**, which is for our example EEG dataset a 1600 × 1600 matrix, in terms of **XX**^T^, which is in our example a 200 × 200 matrix.

For example, in the case of Kronecker shrinkage, equation 5 may be computed as:

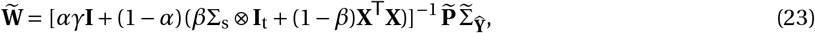

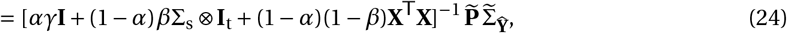

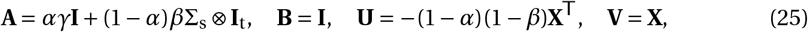

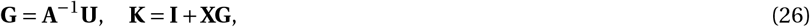

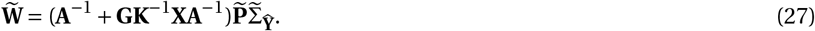

## Appendix C Optimizing the inner cross-validation loop

Our optimization strategy (section 2.10) depends on evaluating the leave-one-out performance of the model many times. The computationally most expensive operation in equation 27 is computing **K**^−1^. However, this matrix only needs to be computed once, whereafter the leave-one-out case where one observation *i* is left out can be obtained efficiently by only computing the change caused by leaving one observation out, instead of re-computing the matrix from scratch. Let **K**_(*i*)_ denote the leave-one-out version of **K**, which in the case of this matrix means the *i*’th row and column are removed. Salmen, Schlipsing, and Igel (2010) have devised an efficient updating algorithm for this case, using the matrix inversion lemma.

Begin by computing **K**_(1)_ and 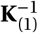 in a conventional manner. Then, **K**_(*i*)_ can be constructed for *i* > 1 by replacing the (*i* − 1)’th row and column of **K**_(1)_ with the first observation. Note that this results in a non-standard ordering of the rows and columns of **K**_(*i*)_, so care must be taken to order the leave-one-out versions of **X** and **Y** in the same manner. The update rule of the inverse can then be formulated as:

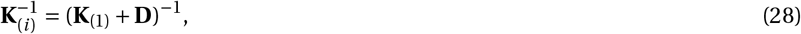

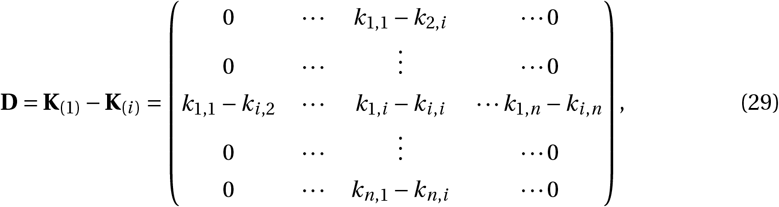

where *k*_*i*,*j*_ refers to the element at row *i* and column *j* of the original matrix **K** and *n* is the total number of observations in **K**.

To apply the inversion lemma (equation 22), **D** must be formulated in terms of **UBV**, which yields:

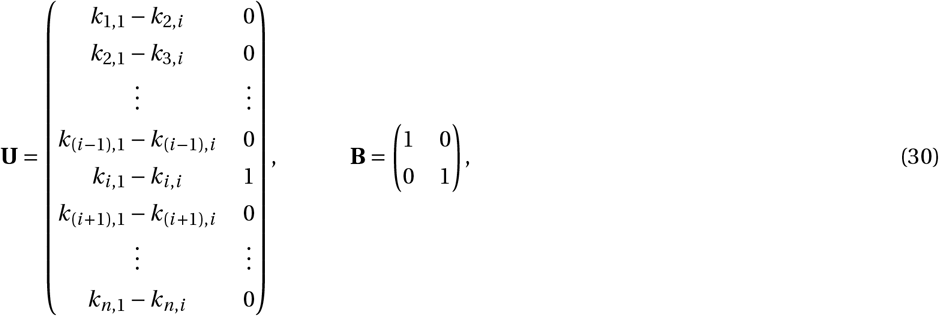

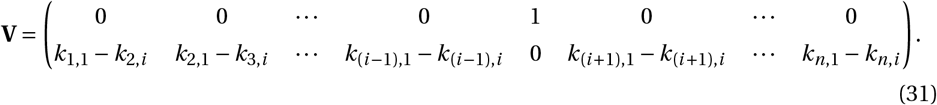

Then, applying equation 22:

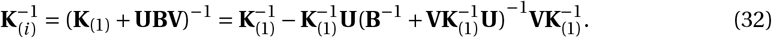

## Footnotes

1 McIntosh and Mišicć, 2013

2 Uusitalo and Ilmoniemi, 1997

3 Jutten and Herault, 1991

4 Vigario, Sarela, Jousmäki, Hamalainen, and Oja, 2000

5 Hämäläinen and Ilmoniemi, 1994

6 Matsuura and Okabe, 1995

7 Van Veen, van Drongelen, Yuchtman, and Suzuki, 1997

8 Gross et al., 2001

9 Grootswagers, Wardle, and Carlson, 2017

10 Lotte, Congedo, Lecuyer, Lamarche, and Arnaldi, 2007

11 Tong and Pratte, 2012

12 Hastie, 2009

13 Pernet, Sajda, and Rousselet, 2011

14 Parra et al., 2003

15 van Vliet et al., 2016

16 Mitchell et al., 2008

17 Huth, Heer, Griffiths, Theunissen, and Jack, 2016

18 Babyak, 2004

19 Blankertz, Lemm, Treder, Haufe, and Müller, 2011

20 Tibshirani, 1996

21 Rifkin and Lippert, 2007

22 Haufe et al., 2014

23 Kohler et al., 1996

24 Lin, Belliveau, Dale, and Hämäläinen, 2006

25 Wipf and Nagarajan, 2009

26 Trujillo-Barreto, Aubert, and Penny, 2008

27 Neely, 1991

28 van Vliet et al., 2014

29 Kutas and Hillyard, 1980

30 Kutas and Federmeier, 2011

31 Hastie, 2009

32 Nelson, McEvoy, and Dennis, 2000

33 van Vliet et al., 2014

34 Gramfort et al., 2013

35 Jas, Engemann, Bekhti, Raimondo, and Gramfort, 2017

36 Perrin, Pernier, Bertrand, and Echallier, 1989

37 Jas et al., 2017

38 Pedregosa et al., 2012

39 van Vliet et al., 2016

40 Babyak, 2004

41 Rifkin and Lippert, 2007

42 Hastie, 2009

43 Rifkin and Lippert, 2007

44 Hastie, 2009

45 Blankertz et al., 2011

46 Engemann and Gramfort, 2015

47 Loan, 2000

48 Bijma, De Munck, and Heethaar, 2005

49 Reuderink, Farquhar, Poel, and Nijholt, 2011

50 Lotte, Guan, and Ang, 2009

51 Fazli et al., 2009

52 Kutas and Hillyard, 1980

53 Kutas and Federmeier, 2011

54 Kutas and Iragui, 1998

55 van Vliet et al., 2016

56 van Vliet, Van Hulle, and Salmelin, 2018

57 van Vliet et al., 2018

58 de Cheveigné and Simon, 2008

59 Byrd, Lu, Nocedal, and Zhu, 1995

60 Pedregosa et al., 2012

61 Kutas and Federmeier, 2011

62 Hämäläinen, Hari, Ilmoniemi, Knuutila, and Lounasmaa, 1993

63 Hämäläinen et al., 1993; Sekihara and Nagarajan, 2008

64 Hämäläinen et al., 1993

65 Kutas and Federmeier, 2011

66 Sekihara and Nagarajan, 2008

67 Blankertz, Tomioka, Lemm, Kawanabe, and Müller, 2008

68 Hoffmann, Vesin, and Ebrahimi, 2006

69 Rivet, Souloumiac, Attina, and Gibert, 2009

70 Delis, Onken, Schyns, Panzeri, and Philiastides, 2016

71 Haufe et al., 2014

72 Parra et al., 2003

73 Kia, Pedregosa, Blumenthal, and Passerini, 2017

74 see electronic supplementary information: “Decoding performance using EOG channels only”

75 Quax, Dijkstra, van Staveren, Bosch, and van Gerven, 2019

76 Treder, Porbadnigk, Shahbazi Avarvand, Müller, and Blankertz, 2016; van Vliet et al., 2016

77 van Vliet et al., 2018

78 Baayen, Davidson, and Bates, 2008

79 Tylavsky and Sohie, 1986

## Notes

#### Summary of Updates

Manuscript underwent a major revision based on the first round of reviewer comments. Refocus of the theoretical framework. Extended Results section with more statistics. Extended Discussion section.

https://aaltoimaginglanguage.github.io/posthoc

